# PALLAS: Penalized mAximum LikeLihood and pArticle Swarms for Inference of Gene Regulatory Networks from Time Series Data

**DOI:** 10.1101/2020.05.13.093674

**Authors:** Yukun Tan, Fernando B. Lima Neto, Ulisses Braga Neto

## Abstract

We present PALLAS, a practical method for gene regulatory network (GRN) inference from time series data, which employs penalized maximum likelihood and particle swarms for optimization. PALLAS is based on the Partially-Observable Boolean Dynamical System (POBDS) model and thus does not require ad-hoc binarization of the data. The penalty in the likelihood is a LASSO regularization term, which encourages the resulting network to be sparse. PALLAS is able to scale to large networks under no prior knowledge, by virtue of a novel continuous-discrete Fish School Search particle swarm algorithm for efficient simultaneous maximization of the penalized likelihood over the discrete space of networks and the continuous space of observational parameters. The performance of PALLAS is demonstrated by a comprehensive set of experiments using synthetic data generated from real and artificial networks, as well as real time series microarray and RNA-seq data, where it is compared to several other well-known methods for gene regulatory network inference. The results show that PALLAS can infer GRNs efficiently and accurately. PALLAS is a fully-fledged program with a commandline user interface, written in python, and available on GitHub (https://github.com/yukuntan92/PALLAS).

## 1 Introduction

Inference of gene regulatory networks (GRN) from gene expression time-series data is a problem of critical importance in Bioinformatics [12]. Many mathematical models have been proposed in the literature to address this problem, including linear models [48, 13], Bayesian networks [37, 18], neural networks [49], differential equations [13, 10] and information theory based approaches [15, 4]. The Boolean network (BN) model [29], is an effective model for GRNs due to its ability to describe temporal patterns of gene activation and inactivation and its comparatively small data requirement for inference [1, 33, 16, 24, 43]. Several extensions of the BN model have been proposed, including Random Boolean Networks [29], Boolean Networks with perturbation (BNp) [41], and Probabilistic Boolean Networks (PBN) [42], and Boolean Control Networks (BCN) [11, 38]. However, all of those models assume that the system Boolean states are completely observable. This is a significant drawback, since all practical methods for the inference of Boolean networks must include a step of ad-hoc binarization of the gene expression data. The Partially-observed Boolean dynamical system (POBDS) model [9] addresses this problem in a principled way, by postulating separate Boolean state and general observation processes. The time-series gene expression data, whether microarray or RNA-seq data, is modeled by the observation process, while the Boolean states are hidden. This allows the optimal inference of the sequence of Boolean states from the time series data, as well as parameter estimation directly from the data.

In this paper, we present PALLAS, a practical method for parametric gene network inference based on the POBDS model, using penalized maximum likelihood and particle swarms for optimization. The penalty in the likelihood score is a Lļ-norm LASSO regularization term [47], which encourages the resulting network to be sparse, i.e., contain a small number of edges between genes; its value can be adjusted by the user to obtain a desired level of sparsity. The likelihood itself is calculated efficiently by an auxiliary particle filter (APF) implementation of the Boolean Kalman Filter [9, 25]. Another novel feature of PALLAS is the application to Boolean models of a particle swarm method: a new mixed continuous-discrete version of the Fish School Search algorithm [6, 7], for efficient simultaneous maximization of the penalized likelihood over the discrete space of networks and the continuous space of observational parameters. An early version of this work appeared previously in a short communication [46]. We mention that particle swarm methods have been employed before for GRN inference using non-Boolean models, namely, recursive neural networks (RNN) in [50] and S-systems in [28].

PALLAS is an extension of the adaptive filtering method proposed in [25]. The latter performs maximization of the likelihood function by exhaustive search over the space of networks and expectation maximization over the space of parameters of the observational model for each candidate network. It is well suited if there is prior knowledge about the network, e.g., most of the edges are known and only a few putative edges are being sought, given the prohibitive computational cost of exhaustively searching the space of all networks. For example, with only 4 genes, there are a total of 688,747,536 Boolean network models to be searched, and with 8 genes this number jumps to approximately 8.8 × 10^32^, rendering exhaustive search completely unfeasible. PALLAS differs from the method in [25] in using penalized maximum likelihood and particle swarms for optimization, which allows it to handle large networks in the absence of any prior knowledge.

The performance of PALLAS is demonstrated by a comprehensive set of experiments. Using synthetic data generated from both real and artificial GRNs, which allows computation of performance metrics, we compare PALLAS to GENIE3 [27], TIGRESS [21], Banjo [44], Best-Fit algorithm [32], REVEAL [34], and GABNI [5]. Using real time series microarray data from the SOS DNA Repair System in E. Coli, we compare PALLAS to the methods in [31, 30, 28]. We also illustrate the performance of PALLAS in recovering known regulatory links in the E. Coli Biofilm Formation Pathway using time series RNA-Seq data.

## 2 Methods

### 2.1 Partially-Observable Boolean Dynamical Systems

The Partially-Observable Boolean dynamical system (POBDS) model [9] allows for uncertainty in Boolean state transitions and partial observation of the Boolean state variables through noise.

#### 2.1.1 State model

Consider a state process {**X**_*k*_; *k* = 0, 1,…}, where **X**_*k*_ ∈ {0, 1}^*d*^ is a Boolean vector of size *d*, which evolves according to

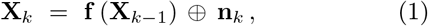

for *k* = 1, 2,… where 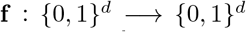 is called the *network function*, **n**_*k*_ ∈ {0, 1}^*d*^ is additive noise at time k, and “⊕” indicates component-wise modulo-2 addition. The state and noise processes are assumed to be independent. The state model (1) can be suitably modified to include external inputs, if desired.

The noise random vector **n**_*k*_ models uncertainty in the state transition: if a component of **n**_*k*_ is 1, the corresponding component of **f** (**X**_*k*−1_) is flipped. As long as all components of **n**_*k*_ have a nonzero probability of being 1, the state process is an ergodic Markov Chain, with a steady state distribution. But if the noise is too intense, i.e., the probability of 1’s in **n**_*k*_ is too large, state evolution becomes chaotic. However, it is well known that important biological pathways are tightly regulated. Accordingly, each component of the noise vector is assumed here to be equal to 1 with a small value *p* = 0.05, independently of the others. A different value 0 ≤ *p* ≤ 0.5 can be selected by the user; however, a value much larger or smaller than 0. 05 is not representative of real gene regulatory networks.

We assume a specific model for the network function that is motivated by gene pathway diagrams commonly encountered in biomedical research. Let a sample state vector **x** ∈ {0, 1}^*d*^ and the network function **f** be expressed in component form as **x** = (*x*_1_,…, *x_d_*) and **f** = (*f*_1_,…, *f_d_*), respectively. Each component *f_i_*: {0, 1}^*d*^ → {0, 1} is given by

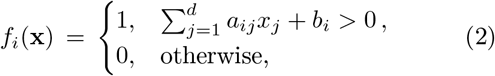

where *a_ij_* = +1 if there is positive regulation (activation) from gene *j* to gene *i*; *a_ij_* = −1 if there is negative regulation (inhibition) from gene *j* to gene *i*; and *a_ij_* = 0 if gene *j* is not an input to gene *i*, whereas *b_i_* = +1/2 if gene *i* is positively biased in the sense that an equal number of activation and inhibition inputs will produce activation; the reverse being the case if *b_i_* = −1/2. The network model is depicted in Figure 1, where the threshold units are step functions that output 1 if the input is positive, and 0, otherwise. This model constraint reduces the number of parameters needed to specify **f** from 2^*d*^ to *d*^2^ + *d*.

**Figure 1:**
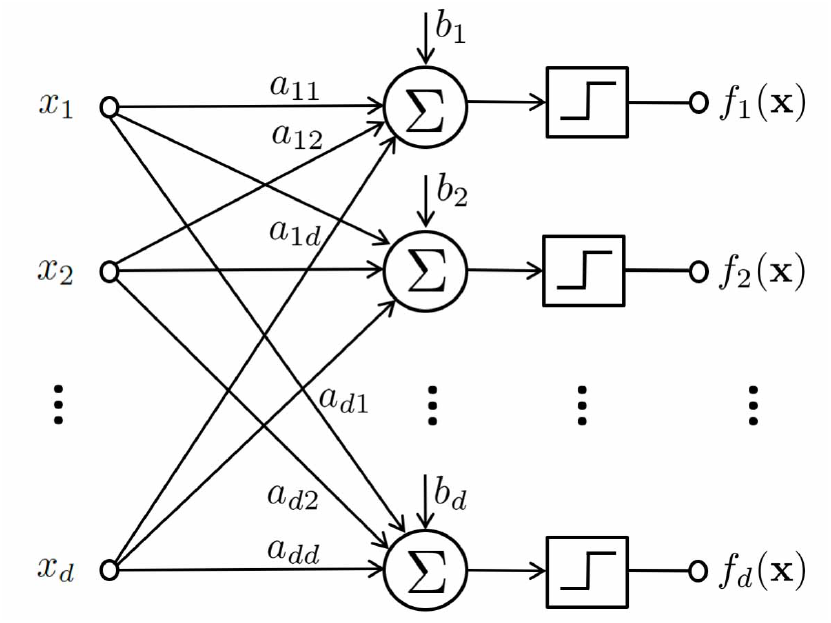
Schematic representation of the network function.

#### 2.1.2 Observation Model

The sequence of states is observed indirectly through the process {**Y**_*k*_; *k* = 0,1,…}, where the measurement vector **Y**_*k*_ is a general nonlinear function of the state and observation noise:

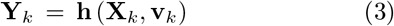

for *k* = 1, 2,…, where the noise vector **v**_*k*_ is assumed to independent of the state process and state transition noise process. We describe next the two observational models considered in this paper, corresponding to two common gene expression modalities: RNA-Seq count data or microarray fluorescence data. Observational models for other data modalities can be introduced, if desired.

##### RNA-Seq observation model

RNA-Seq data can be modeled with the Poisson distribution [19] or the negative binomial distribution [20, 2]. Here, we employ the latter, since it is able to address overdispersion in the count distributions. We assume that the transcript counts **Y**_*k*_ = (*Y*_*k*1_,…, *Y_kd_*) are re-lated to the state **X**_*k*_ = (*X*_*k*1_,…, *X_k_d*) via

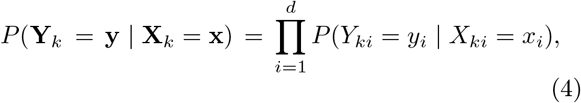

and adopt the negative binomial model for each count,

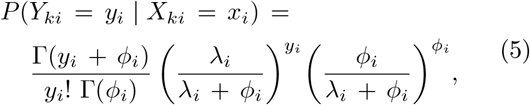

where Γ denotes the Gamma function, and *ϕ_i_*, λ_*i*_ > 0 are the real-valued inverse dispersion parameter and mean read count of transcript *i*, respectively, for *i* = 1,…, *d*. The inverse dispersion parameter *ϕ_i_* specifies the amount of observation noise: the larger it is, the less observation noise is present. We model the parameter λ_*i*_ in log-space as:

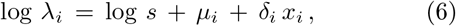

where the parameter *s* is the sequencing depth, which depends on the instrument, *μ_i_* ≥ 0 is the baseline level of expression in the inactivated transcriptional state, and *δ_i_* > 0 is the difference between read count as gene *i* goes from the inactivated (*x_i_* = 0) to the activated (*x_i_* = 1) state, for *i* = 1,…, *d*.

##### Microarray observation model

A reasonable model for continuous microarray fluorescence data is a Gaussian linear model:

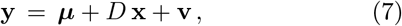

where ***μ*** = (*μ*_1_,…, *μ_d_*) ≥ 0 is the vector of baseline expression levels corresponding to the “zero” or inactive state for each gene, *D* = diag{*δ*_1_,…, *δ_d_*} > 0 is a diagonal matrix containing differential expression values for each gene, and 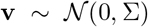 is an uncorrelated zero-mean Gaussian noise vector, where 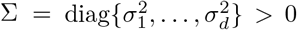. Notice that (4) is still satisfied here.

#### 2.1.3 Boolean Kalman Filter

Given a time series of observations **Y**_1:*k*_ = {**Y**_1_,…,**Y**_*k*_}, the Boolean Kalman Filter[9] (described in detail in the Supplementary Material) computes exactly the minimum mean-square error state estimator:

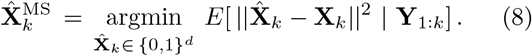

The BKF also computes the probabilities needed to determine the likelohood function, as detailed in Section 2.2.1. When the network is large, however, computation of the BKF is intractable since each transition matrix contain 2^2*d*^ elements which requires large computation and memory. In this case, approximate methods must be used, such as the Sequential Monte Carlo (SMC) method, also known as particle filter [3]. Here we use the auxiliary particle filter implementation of the Boolean Kalman Filter (APF-BKF), described in [26] (please see that reference for the details).

### 2.2 PALLAS Algorithm

In this section, we describe in detail PALLAS (Penalized mAximum LikeLihood and pArticle Swarms), an algorithm for inference of Boolean gene regulatory networks from noisy time series of gene expression data. The algorithm has two main components: 1) efficient computation of a penalized log-likelihood cost function; 2) maximization of the previous cost function using a novel particle swarm method, namely, a mixed discrete-continuous fish school search procedure. We describe here the general case, where no prior knowledge is available to set model parameters. The algorithm can be easily modified to allow some of the parameters to be specified by the user, by simply reducing the size of the parameter space and using the known parameter values in the likelihood computation.

Let *θ* = (*θ*_disc_, *θ*_cont_) ∈ Θ> with *θ*_disc_ ∈ Θ_disc_ and *θ*_cont_ ∈ Θ_cont_, be the discrete and continuous unknown model parameters, where Θ, Θ_disc_ and Θ_cont_ are the corresponding parameter spaces, with Θ = Θ_disc_ × Θ_cont_. Here, *θ*_disc_ contains the parameters of the network function in (2), namely the edge parameters *a_ij_* ∈ {−1, 0, 1}, for *i, j* = 1,…, *d*, and the regulation bias parameters *b_i_* ∈ {-1/2, 1/2}, for *i* = 1,…, *d*. Hence, Θ_disc_ = { −1, 0, 1}^*d*^2^^ × {−1/2, 1/2}^*d*^. This is a finite space, but its cardinality |Θ_disc_| = 3^*d*^2^^ × *2^*d*^* increases extremely fast with the number of genes *d*. For example, for a network with only *d* = 4 genes, |Θ_disc_| = 688747536, while if *d* = 8, then |Θ_disc_| ≈ 8.8 × 10^32^. On the other hand, *θ*_cont_ contains the observational parameters: the baseline expression levels *μ_i_* > 0 and the differential expression levels *δ_i_* > 0, for *i* = 1,…, d, for both RNA-Seq and microarray data, the inverse dispersion parameters *ϕ_i_* > 0, for *i* = 1,…, *d*, for RNA-Seq data, and the standard deviations *σ_i_* > 0, for *i* = 1,…, d, for microarray data (the sequencing depth parameter s is assumed known for a given RNA-seq assay, so it is not part of *θ*_cont_). Hence, the dimensionality of *θ*_cont_ is *Q* = 3*d* in both cases.

The mixed discrete-continuous fish school search procedure employed by PALLAS assumes that the parameter space is a closed and bounded region with an absorbing decision boundary (if the current best estimate exceeds the boundary, it remains at the boundary). This is not a limiting requirement in practice, since sensible lower and upper bounds can be set for all the observational parameters. These intervals can be set by the user, or the following data-driven procedure to obtain default intervals is employed. Let min, max, and mean be respectively the minimum, maximum, and mean value of the observed data for all genes across all time points and available time series. In the case of RNA-Seq data, the data must be normalized by dividing the measurements by the sequencing depth and then taking logs prior to computing the mean, max, and mean values. Then the following intervals are assumed:

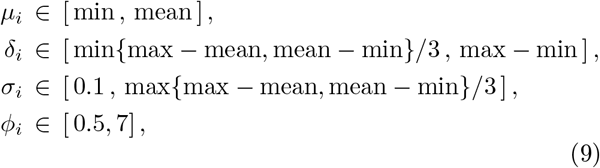

for *i* = 1,…, *d*. Some of the parameters can often be assumed to be the same across different genes, which reduces the data requirement of the estimation problem. In the simplest case, there are single parameters *μ, δ*, and *σ* or *ϕ* for all genes, so that *Q* = 3. In any case, Θ_cont_ is a closed and bounded rectangular region in 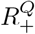.

Next we describe the two main steps comprising the PALLAS algorithm: computation of the penalized log-likelihood function and the novel mixed discrete-continuous fish-school search method.

#### 2.2.1 Penalized Maximum-Likelihood Computation

Suppose that the sample data consist of *n* independent time series 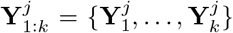 up to time *k*, for *j* = 1,…, *n*. The penalized log-likelihood of model *θ* at time *k* is defined as

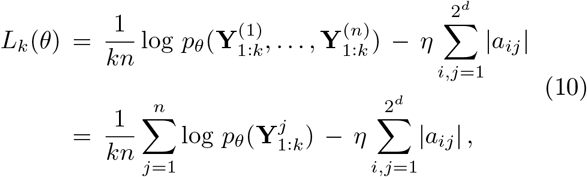

where *η* > 0 is a regularization parameter, which has a default value of *η* = 0.01 in our implementation. Hence, the penalized log-likelihood in (10) is the sum of the average log-likelihood per time series and a negative value times the number of edges in the model. Maximization of (10) thus encourages the model to both fit the data and be sparse, i.e., contain a small number of edges between genes, which is in agreement with biological knowledge. The value of η can be adjusted by the user to obtain a desired level of sparsity. Notice that

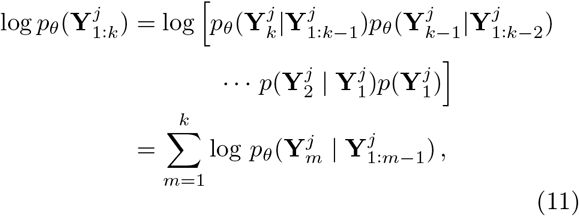

where

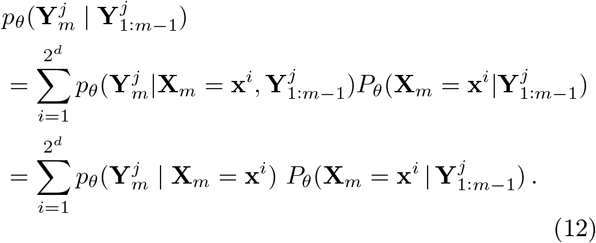

With 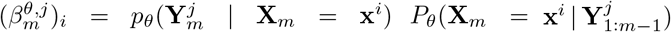, the penalized log-likelihood in (10) be written as

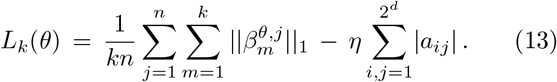

The sequence of values 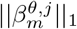, for *j* = 1,…,*n* and *m* = 1,…, *k*, can be computed by a BKF tuned to parameter *θ* applied to the time series 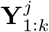 (see the Supplementary Material for a description of the BKF). As mentioned in the previous section, here we use the auxiliary particle filtering implementation of the BKF, for computational efficiency. The maximum-likelihood estimator of parameter *θ* at time *k* is then given by

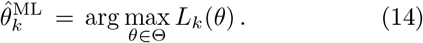

A state estimate 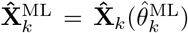 can be obtained, if desired, where 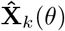 denotes the optimal state estimator produced by a BKF tuned to the parameter *θ*.

#### 2.2.2 Mixed Fish School Search Algorithm

In this section, we describe in detail a novel particleswarm algorithm for optimization over a combined discrete-continuous parameter space, called the mixed fish school search (MFSS) algorithm, which is an extension of the fish school search (FSS) algorithm for continuous parameter spaces proposed in [6].

In the MFSS algorithm, the objective is to find a model that maximizes a given score or fitness — in our present case, this is the penalized log-likelihood defined in the previous section. Each candidate model, i.e., each candidate parameter vector *θ* = (*θ*_disc_, *θ*_cont_), corresponds to a particle or “fish.” The length of *θ* is denoted by P. From the previous section, *P* = *d*^2^ + *d* + *Q*. The fish school is an ensemble of *S* such particles in the parameter space Θ = Θ_disc_ ×Θ_cont_. The position of fish *s* at iteration *r* will be denoted by 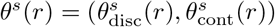, for *s* = 1,…, *S*, and *r* = 0,…, *R*. The number of fishes S and the total number of iterations *R* are user-defined parameters (in practice, *S* = 3 × *P* and *R* = 5000 are found to be good values). In addition, each fish *s* has a weight *w_s_*(*r*) at iteration *r*, which reflects the quality of the solution.

##### Initialization

The initial position 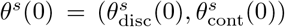 of each fish is assigned randomly. The continuous vector 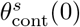 is drawn from a uniform distribution over Θ_cont_, but for the discrete part, it is advantageous to use a non-uniform distribution to initialize the edge parameters, in such a way that 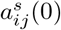 is equal to −1 or 1 with probability 1/4, and 0 with probability 1/2, for *i,j* = 1,…, *d*, which introduces a bias towards 0 over 1 and −1. This is in agreement with the biological observation that GRNs tend to be sparsely connected. The initial value 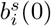 (0) of the regulation bias parameter is chosen to be either −1/2 or 1/2 with equal probabilities, for *i* = 1,…, *d*.

##### Individual movement operator

This is an exploratory step, where each fish independently moves a short distance in a random direction, as long as this increases the fitness function. Let 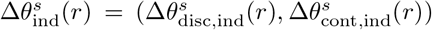 be the (candidate) individual displacement vector for fish *s* at iteration *r*. Vector 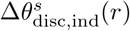 is drawn from a uniform distribution over the rectangular region [−1, 1]^*d*^2^+*d*^, while 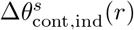 is drawn from a uniform distribution over the rectangular region [−*τ*_1_(*r*),*τ*_1_(*r*)] × ⋯ × [−*τ_Q_*(*r*), *τ_Q_*(*r*)]. The step size bounds *τ_q_*(*r*), for *q* = 1,…, *Q*, shrink linearly with *r*, in order to ensure convergence and emphasize exploitation over exploration at later iterations. In our implementation, the initial and final values *τ_q_* (1) and *τ_q_*(*R*) are set, respectively, to 10% and 0.01% of the range (i.e., the difference between upper and lower bounds) of the corresponding continuous parameter — these values can be modified by the user, if desired. Now, 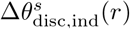 needs to be quantized into the lattice {−1,0,1}^*d*^2^+*d*^ in order to be added to the discrete component of the current fish position. The quantization scheme we adopt here is a generalization of the method for binary parameters in [40]. We define two adaptive thresholds:

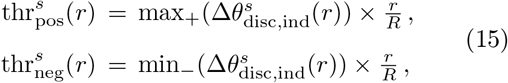

where the operator max_+_(**v**) is equal to the maximum of the components of vector **v** if at least one of them is positive, and equal to zero, otherwise; similarly, min_(**v**) is equal to the minimum of the components of **v** if at least one of them is negative, and equal to zero, otherwise. The factor r/R increases the thresholds (in magnitude) with r, to favor exploitation over exploration at later iterations and ensure convergence.

The quantized discrete displacement vector is obtained by assigning 1 to a positive component if it is larger than 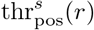, assigning −1 to a negative component if it is smaller than 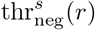, and assigning 0 to all other components (no movement). Then the position of fish s is updated if the exploratory move causes an increase in fitness:

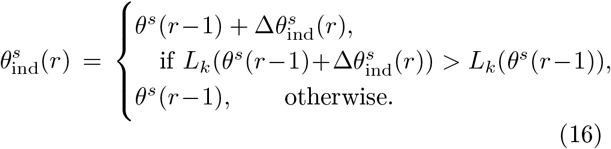

where *L_*k*_* is the penalized log-likelihood of the model, defined in the previous section. An absorbing boundary condition is adopted, whereby each fish interrupts its movement at the boundary of the parameter space, at the point where it encounters it.

##### Feeding operator

The weights of all fish are updated based on the fitness improvement from the previous individual movement, if any:

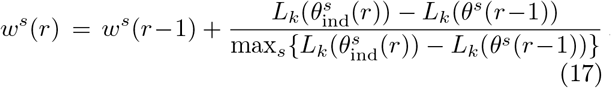

##### Collective instinctive movement operator

This operator makes the fish that had successful individual movements influence the collective direction of movement of the school. The position of each fish s is updated according to:

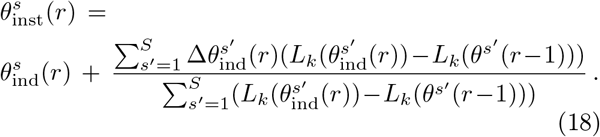

The displacement in discrete parameter space is quantized following the same procedure adopted to discretize the individual movement displacement vector.

##### Collective volitive movement operator

This is similar to the individual movement step, but now the fish move in concert, depending on whether the fish school is successful after the previous steps, i.e., its total weight increases, or not. If the fish school is successful, then it should contract, changing from exploration to exploitation mode. Otherwise, it should expand in order to explore the space more. This is accomplished by first defining the current fish school barycenter:

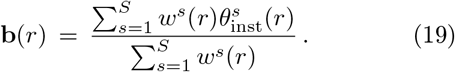

For each fish *s*, after the collective instinctive movement at iteration *r*, let 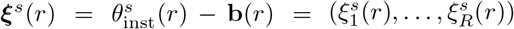 be the position vector with respect to the school barycenter. Let 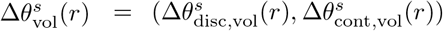 be the collective volitive displacement vector for fish *s* at iteration *r*. Vector 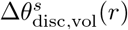 is drawn from a uniform distribution over the rectangular region 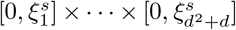 and quantized by the same process used in the individual move, while 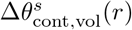 is drawn from uniform distribution over the rectangular region 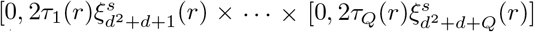, where *τ*_1_(*r*),…,*τ_Q_*(*r*) are the same step sizes used in the individual movement step. If the school is successful, i.e., if 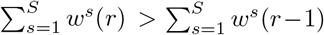, then its radius should contract, and

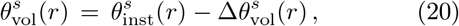

otherwise, the radius expands, so the school can escape a bad region, and

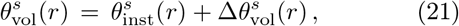

The result is the new position of the fish *θ^s^*(*r*).

## 3 Results

In this section, we present the result of a comprehensive set of numerical experiments, using both synthetic and real gene expression time series data, to assess the performance of PALLAS and compare it against that of other popular methods in the literature. No prior knowledge is used; i.e., all model parameters must be estimated. Unless otherwise noted, the default values for all PALLAS fixed parameters and estimation intervals are used, as described in the previous sections.

### 3.1 Performance Criteria

The problem of comparing networks is a nontrivial one; there is not a single metric that captures both the topological and dynamical properties of the networks [14]. Here we consider two classes of metrics, one based on the difference between the network functions, which takes into account the full regulatory relationships among genes, and the other based on edge-calling error rates, which considers only the network topology.

#### 3.1.1 Network Function Distance

Let **f** = (*f*_1_,…, *f_d_*) and 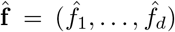 be the network functions of the groundtruth and inferred networks, where the component functions *f_i_* and 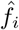 are Boolean functions on *d* variables, for *i* = 1,…, *d*; see (1). The performance criterion is the average number of disagreeing Boolean functions between the two networks

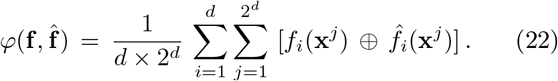

This distance is related to the dynamical behavior of the networks, since it has to do with how the Boolean functions differ.

#### 3.1.2 Edge-Calling Error Rates

An *edge* in the groundtruth network represents a relationship between two genes. Here we consider directionality (an edge from gene *i* to gene *j* is distinct from an edge from gene *j* to gene *i*), but disregard activation/inhibition relationships (this is done because some of the methods to which PALLAS is compared in this section do not capture activation/inhibition). Let TP and FN be the total number of directional edges that are correctly detected (irrespective of inhibition/activation) and incorrectly missed by the inference algorithm, respectively. Similarly, let FP and TN be the total number of directional edges that are incorrectly found and correctly missed, respectively. We define the following edge-calling error rates:

a. Sensitivity/True Positive Rate (TPR):

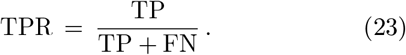
b. Specificity/True Negative Rate (SPC):

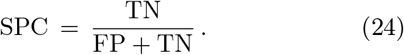
c. Precision/Positive Predictive Value (PPV):

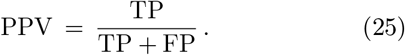

### 3.2 Experiments with Synthetic Data

#### 3.2.1 Mammalian Cell-Cycle Network with Synthetic RNA-Seq Data

Here, we present results based on the well-known Mammalian Cell-Cycle network [17], which is displayed in Figure 2. (Results for a different GRN are presented in the Supplementary Material). The state vector is **X** = (CycD, Rb, p27, E2F, CycE, CycA, Cdc20, Cdh1, UbcH10, CycB). This is a large network, with a huge parameter space, for which the estimation problem is hard. The gene interaction parameters *a_ij_* can be read from Figure 2 in the same way as in the last section. Once again, the regulation biases are set to bi = −1/2, for *i* = 1,…, 10. The transition noise parameter *p* is selected randomly in the interval [0.01,0.1]. The RNA-Seq data model parameters are *μ_i_* ≡ *μ* = 0.1, *δ_i_* ≡ *δ* = 3, *ϕ_i_* ≡ *ϕ* = 5, for *i* = 1,…, 10. The sequencing depth is set to *s* = 22.52 (500K-550K reads) and the time series length is fixed at 5O. Here we compare PALLAS with the GENIE3 [27], TIGRESS [21], and Banjo [44] algorithms. Like PALLAS, these algorithms can operate directly on the noisy time series, without a need for ad-hoc binarization. However, they do not estimate observational parameters or provide activation/inhibition information, so only the edge-calling error rates in Section 3.1.2 are appropriate here. Average rates obtained over 20 repetitions of the experiment are displayed in Figure 3. One can see that with similar specificity, PALLAS displays higher sensitivity and precision than GENIE3 and TIGRESS. Although it was not possible to adjust the specificity of Banjo to the same levels, we can see that its sensitivity is quite low. In fact, Banjo returned a very small number of edges overall in this experiment. PALLAS also displayed the highest precision among all the algorithms.

**Figure 2:**
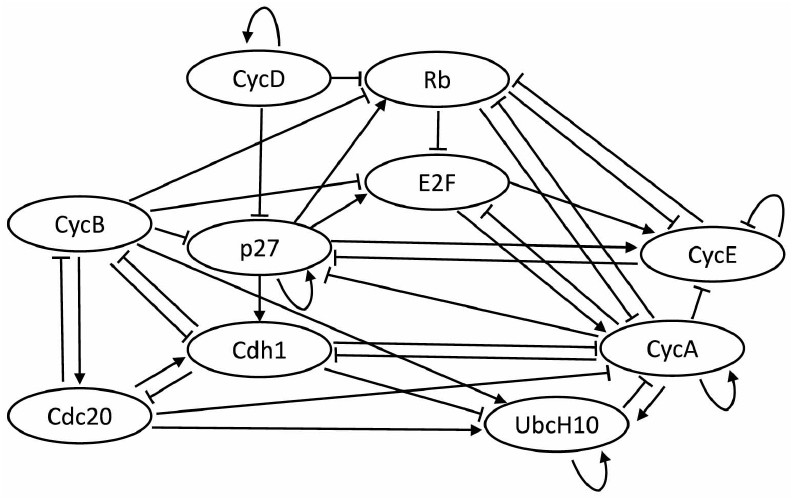
Mammalian cell cycle network.

**Figure 3:**
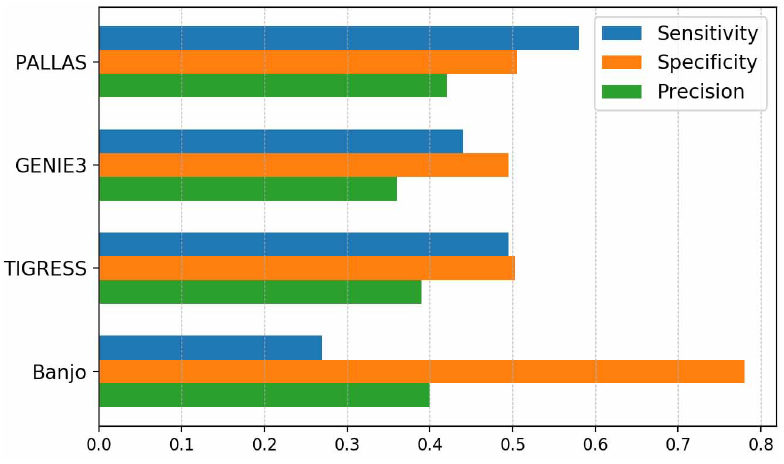
Mammalian cell cycle experiment results.

#### 3.2.2 Artificial Networks with Synthetic RNA-Seq Data

In this section we report results obtained on an ensemble of 10 randomly generated networks with d = 8 genes, where each gene is regulated by 3 other genes on average. Edge connectivity, including activation and inhibition, as well as regulation biases, are randomly chosen. The transition noise parameter *p* is selected randomly in the interval [0.01,0.1]. RNA-Seq synthetic data are generated with parameters *μ_i_* ≡ *μ* = 0, *ϕ_i_* ≡ *ϕ* = 1 or 5, for *i* = 1,…, 8. In the first case, there is more observation noise, and the problem is harder. The parameters *δ_i_* are allowed to vary uniformly over the intervals [1, 2] or [1, 5], for *i* = 1,…, 8. In the first case, the problem is harder, since the differences in observed expression are smaller. Sequencing depth is set at s = 22.52 (500K-550K reads).

Here, we compare PALLAS with the Best-Fit [32], REVEAL [34], and GABNI [5] algorithms. These methods apply to Boolean time series, so they need to employ ad-hoc binarization of the gene expression data. For the first two, [8] recommends the use of the KM3 binarization method, while for GABNI, [4] recommends the use of K-means binarization; hence, we use those binarization methods here. The output of the Best-Fit and REVEAL algorithms are Boolean transition functions, for which the network function distance is appropriate. On the other hand, the output of GABNI consist of positive (activating) or negative (inhibitory) interactions, for which we use the edge-calling error rates defined previously.

Average network function distances and edge-calling error rates obtained over 20 repetitions of the experiment (2 for each of the 10 networks) with *ϕ* = 5 are displayed in Figures 4 and 5 (corresponding results for *ϕ* = 1 are shown in the Supplementary Material). Figure 4 shows that the performance of Best-Fit and PALLAS increases with the time series length, while the performance of REVEAL is mostly stable. PALLAS perform better than the Best-Fit algorithm, especially when δ is smaller. This reflects the fact that ad-hoc binarization of the data becomes less accurate with a smaller difference between activation/inactivation levels in the observed data, which is determined by δ. Figure 5 shows that PALLAS beats GABNI in sensitivity throughout, as well as in specificity under low observation noise and sufficient data. Indeed, the specificity of GABNI is artificially large for small amounts of data, when it detects very few edges.

**Figure 4:**
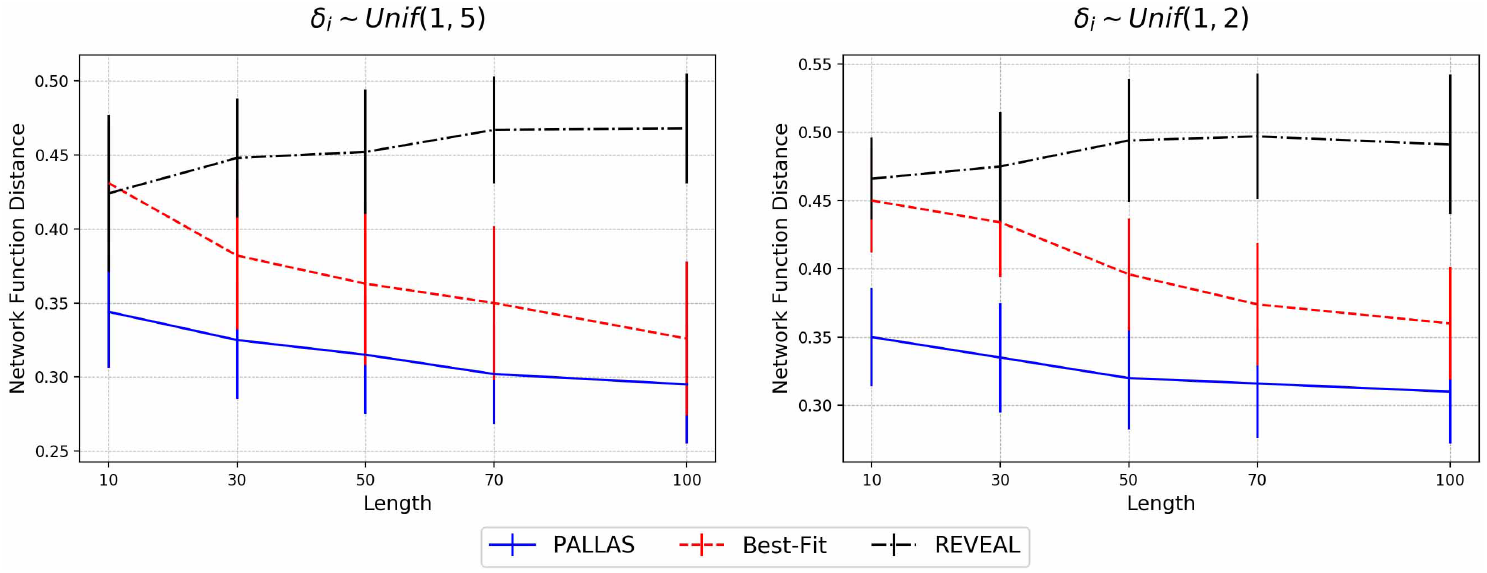
Comparison of network function distance among the PALLAS, Best-Fit, and REVEAL algorithms, under different *δ* ranges.

**Figure 5:**
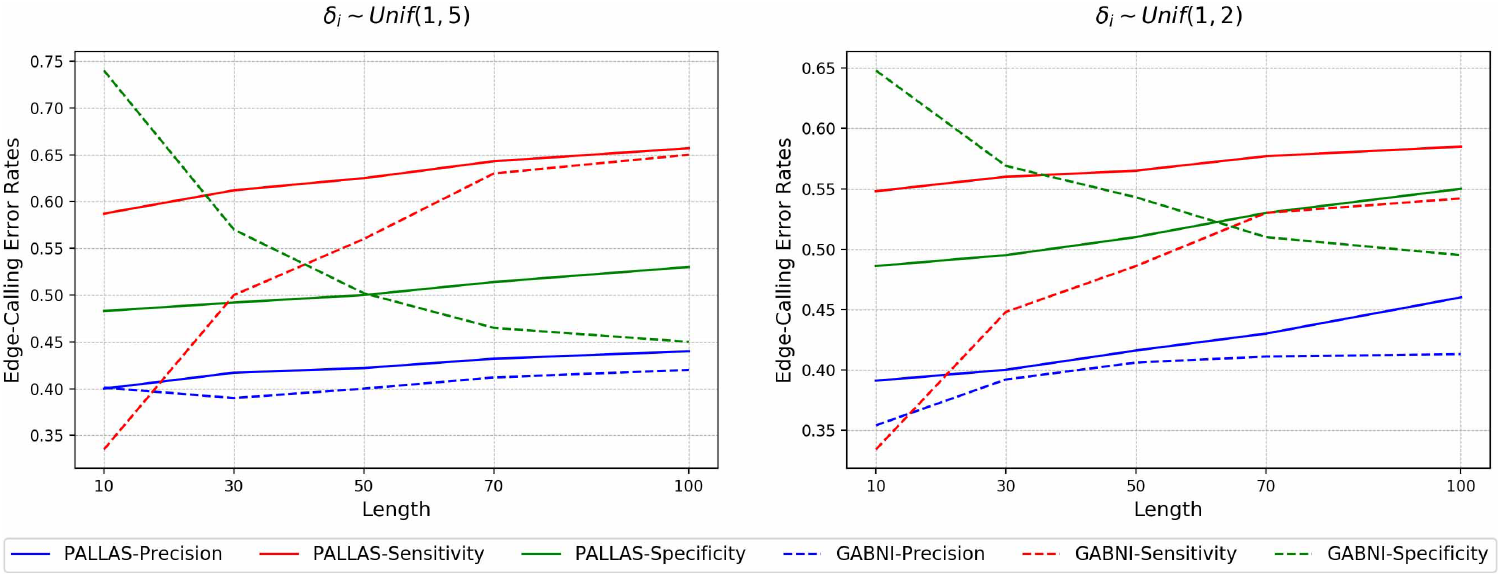
Comparison of edge-calling error rates between the PALLAS and GABNI algorithms, under different *δ* ranges.

### 3.3 Experiments with Real Data

#### 3.3.1 E. Coli SOS DNA Repair System

In this section, we demonstrate the application of PALLAS to real microarray data from a well-known biological system, namely, the SOS DNA repair system in E. Coli. In the normal state, the protein *LexA* is known to be a repressor to the SOS genes. When DNA is damaged, the protein *RecA* becomes activated and mediates *LexA* autocleavage, which causes activation of the SOS genes. After the activated SOS genes repair the damaged DNA, *RecA* stops mediating *LexA* autocleavage and *LexA* represses the SOS genes again. The full SOS DNA repair gene network is displayed in Figure 6 [45, 28]. We attempt to infer this network from gene expression time series datasets generated by [39] (http://www.weizmann.ac.il/mcb/UriAlon/download/downloadable-data). Each time series contains 50 measurements for every 6 minutes including the initial zero concentrations; we pick the third dataset in the database for this experiment, and compare the results against those found in [31, 30, 28].

**Figure 6:**
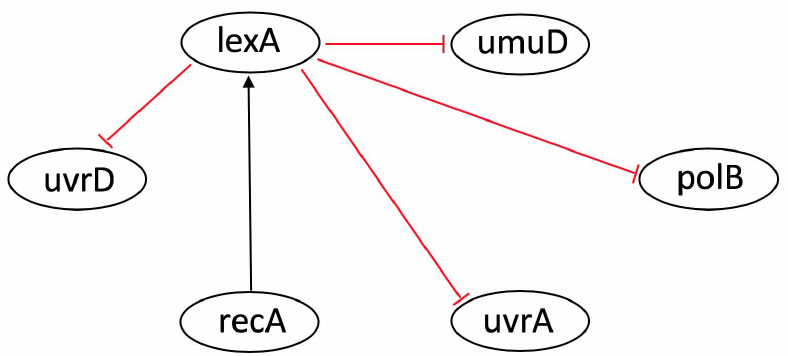
SOS DNA repair system in E.coli (the red edges are the ones successfully recovered by PALLAS).

The sparsity parameter λ in (10) is chosen to produce about half of the possible edges in the six-gene network. Figure 6 displays in red the edges of the original network that were successfully recovered by a consensus of the top three networks found by PALLAS, according to the penalized likelihood score (the full network is displayed in the Supplementary Material). We can see that all inhibitory edges from *lexA* were successfully detected. Although PALLAS infers the wrong direction between *recA* and *lexA*, the connection is detected. With a similar total number of inferred edges, [30] finds the opposite regulations, i.e., all the inhibitory edges are inferred as activating edges. While [31] finds most of the inhibitory edges, it misses the important edge from *lexA* to *uvrA.* Finally, [28] recovers only two of the edges.

#### 3.3.2 E. Coli Biofilm Formation Pathway

In this section, we demonstrate the performance of PALLAS on RNA-Seq time series expression data from a pathway involved in biofilm formation by E. Coli, namely, the Rpos(sigmaS)/MlrA/CsgD cascade, which involves eight genes: *Rpos, MlrA, CsgD, YciR, YoaD, BcsA, YaiC, YdaM*. Information on this pathway can be found in the KEGG database (https://www.genome.jp/kegg/) as well as in [22, 35, 36]. Figure 7 displays a consensus gene network derived from these sources. The gene expression data used is from the E. Coli Strain B/REL606 and is available at the Dryad Digital Repository (https://datadryad.org/resource/doi:10.5061/dryad.hj6mr) [23]. This dataset consists of 3 bacterial samples and 9 time points evenly spaced for each sample. The genes in this pathway display similar values at low expression levels, but vary considerably at high expression levels. Accordingly, we assume a single baseline parameter *μ_i_* ≡ *μ* for all genes, but the parameters *δ_i_* and *ϕ_i_* are allowed to differ from gene to gene, for *i* = 1,…, 8. The sequencing depth is set at s = 1.02 (1k-50k reads) reflecting the low read counts in the data set.

**Figure 7:**
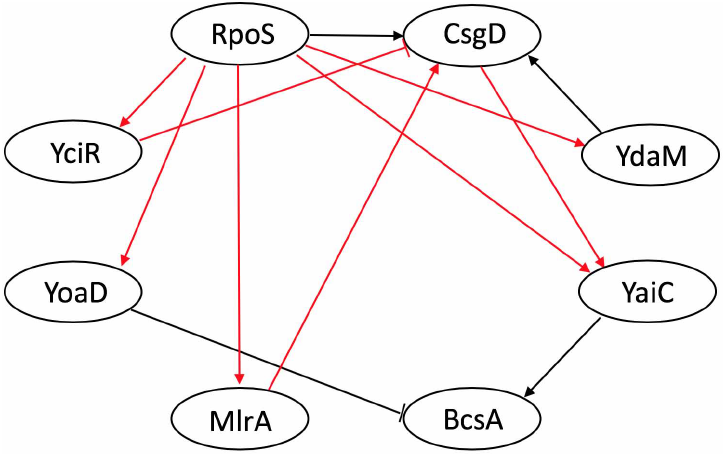
Biofilm architecture of Escherichia coli (the red edges are the ones successfully recovered by PALLAS).

As in the previous experiment, the sparsity parameter λ in (10) is chosen to produce about half of the possible edges in the eight-gene network. Figure 7 displays in red the edges of the original network that were successfully recovered by a consensus of the top three networks found by PALLAS, according to penalized likelihood score (the full network is displayed in the Supplementary Material). PALLAS successfully infers five out of the six important activating interactions from *RpoS*. Most of the other connections in the original network were correctly detected.

## 4 Conclusion

We presented in this paper PALLAS, a new framework for inference of Boolean gene regulatory networks from gene expression time series data. The algorithm avoids ad-hoc binarization of the gene expression data and allows inference of large networks by employing penalized maximum likelihood as a regularization method, applying particle filtering for the computation of the likelihood, and using a novel version of the fish school search particle swarm algorithm to search the parameter space. Numerical experiments using synthetic time series data show that PALLAS outperforms other well-known GRN inference methods. The performance of PALLAS was also demonstrated on real gene expression time series data from the SOS DNA repair and Biofilm formation pathways in E. Coli.

## Notes

https://www.researchgate.net/publication/338685898

### Competing Interest Statement

The authors have declared no competing interest.

