## Supplementary Material for "PALLAS: Penalized mAximum LikeLihood and pArticle Swarms for Inference of Gene Regulatory Networks from Time Series Data"

### 1 Boolean Kalman Filter

For a vector $\mathbf{u} \in [0,1]^d$, define the threshold operator $\overline{\mathbf{u}} \in \{0,1\}^d$ as $\overline{\mathbf{u}}(i) = 1$ if $\mathbf{u}(i) > 1/2$ and 0 otherwise, for $i = 1, \ldots, d$, respectively. It was shown in [Imani and Braga-Neto(2017)] that

$$\hat{\mathbf{X}}_k^{\mathrm{MS}} \,=\, \overline{E\left[\mathbf{X}_k \mid \mathbf{Y}_{1:k}\right]}\,. \tag{1}$$

Assuming that all the parameters of the state and observation models *are known*, then the optimal MMSE filter in (1) can be calculated exactly by a recursive procedure called the Boolean Kalman filter (BKF) [Braga-Neto(2011)], which is described briefly next.

Let $(\mathbf{x}^1, \ldots, \mathbf{x}^{2^d})$ be an arbitrary enumeration of all state vectors. Define the state conditional probability distribution vector $\mathbf{\Pi}_{k|k}$ with components

$$(\mathbf{\Pi}_{k|k})_i \,=\, P\left(\mathbf{X}_k = \mathbf{x}^i \mid \mathbf{Y}_{1:k}\right)\,,\;\; i = 1, \ldots, 2^d, \tag{2}$$

for $k = 0, 1, \ldots$ According to equation (1),

$$\hat{\mathbf{X}}_k^{\mathrm{MS}} \,=\, \overline{E\left[\mathbf{X}_k \mid \mathbf{Y}_{1:k}\right]} \,=\, \overline{A\mathbf{\Pi}_{k|k}}\,,\;\; k = 1, 2, \ldots, \tag{3}$$

where $A = \left[\mathbf{x}^1 \cdots \mathbf{x}^{2^d}\right]$ is a matrix of size $d \times 2^d$.

The computation of $\mathbf{\Pi}_{k|k}$ can be performed recursively. First, we have

$$\mathbf{\Pi}_{k|k-1} \,=\, M_k \mathbf{\Pi}_{k-1|k-1}\,,\;\; k = 1, 2, \ldots \tag{4}$$

where $M_k$ is the *transition matrix* of the Markov state process, with entries

$$(M_k)_{ij} \,=\, P(\mathbf{X}_k = \mathbf{x}^i \mid \mathbf{X}_{k-1} = \mathbf{x}^j) \,=\, P(\mathbf{n}_k \,=\, \mathbf{x}^i \oplus \mathbf{f}(\mathbf{x}^j)),\;\;\; i,j = 1, \ldots, 2^d. \tag{5}$$

On the other hand,

$$\mathbf{\Pi}_{k|k} \,\propto\, T(\mathbf{Y}_k)\,\mathbf{\Pi}_{k|k-1}\,,\;\; k = 1, 2, \ldots \tag{6}$$

where "$\propto$" means that the result must be normalized to add up to 1, and the *update matrix* $T(\mathbf{Y}_k)$ is diagonal of size $2^d \times 2^d$, with diagonal elements:

$$(T_k(\mathbf{Y}_k))_{ii} \,=\, p\left(\mathbf{Y}_k \mid \mathbf{X}_k = \mathbf{x}^i\right),\;\; i = 1, \ldots, 2^d\,. \tag{7}$$

According to equations (4)–(6) in the main text, the update matrix in (7) for the RNA-seq model is given by:

$$\begin{aligned}(T_k(\mathbf{y}))_{ii} \,&=\, P(\mathbf{Y}_k \,=\, \mathbf{y} \mid \mathbf{X}_k \,=\, \mathbf{x}^i)\\ &=\prod_{j=1}^{d}\left[\frac{\Gamma(y_j+\phi_j)}{y_j!\;\Gamma(\phi_j)}\left(\frac{s\,\exp(\mu_j+\delta_j\,x_j^i)}{s\,\exp(\mu_j+\delta_j\,x_j^i)+\phi_j}\right)^{y_j}\left(\frac{\phi_j}{s\,\exp(\mu_j+\delta_j\,x_j^i)+\phi_j}\right)^{\phi_j}\right],\end{aligned} \tag{8}$$

for $i = 1, \ldots, 2^d$. For the microarray model, it follows from equation (7) in the main text that the update matrix is given by:

$$(T_k(\mathbf{y}))_{ii} = P(\mathbf{Y}_k = \mathbf{y} \mid \mathbf{X}_k = \mathbf{x}^i) = \frac{1}{(2\pi)^{\frac{d}{2}} \prod_{j=1}^{d} \sigma_j} \times \exp\left(-\frac{1}{2}\sum_{j=1}^{d}\left(\frac{y_j - \mu_j - \delta_j\, x_j^i}{\sigma_j}\right)^2\right), \quad (9)$$

for $i = 1, \ldots, 2^d$.

#### 2 Random Networks with Synthetic RNA-Seq Data

Average network function distances and edge-calling error rates obtained over 20 repetitions of the experiment (2 for each of the 10 networks) with $\phi = 1$ are displayed in Figures 1 and 2.

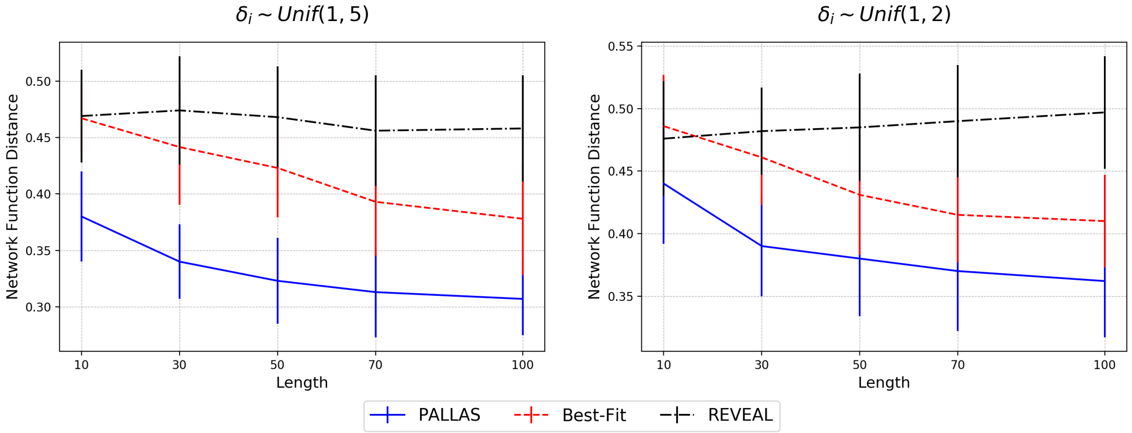


Figure 1: Comparison of network function distance for the three algorithms with different $\delta$ ranges.

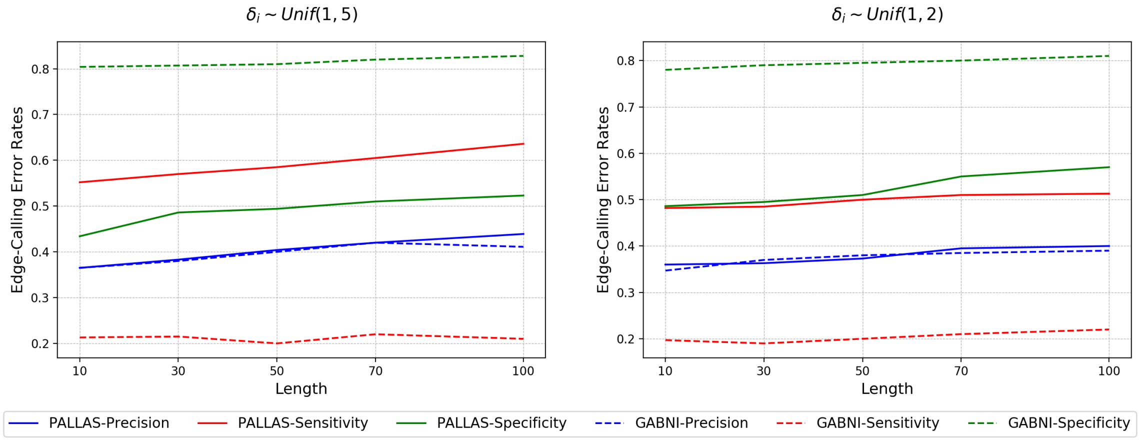


Figure 2: Comparison of edge-calling error rates for the three algorithms with different $\delta$ range.s

### 3 p53-MDM2 Negative-Feedback Loop Gene Regulatory Network with Microarray Data

The experiments in this section use the well-known p53-MDM2 negative-feedback gene regulatory network [Batchelor *et al.*(2009)], which is displayed in Figure 3. The state vector is $\mathbf{X}$ = (ATM, p53, Wip1, MDM2), while dna_dsb acts an external Boolean input that signals DNA damage (p53 is a master tumor-suppressing gene that activates DNA repair mechanisms). The gene interaction parameters $a_{ij}$ can be read from Figure 3.

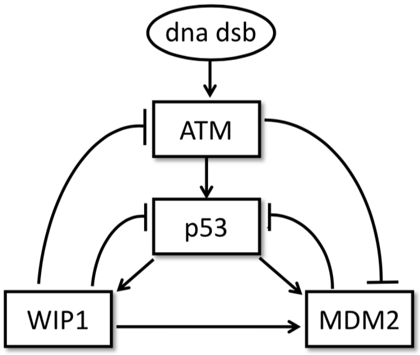


Figure 3: p53-MDM2 negative-feedback loop gene regulatory network.

For example, p53 is activated by ATM, and is inhibited by WIP1 and MDM2. These interactions can be represented as $a_{21} = +1, a_{22} = 0, a_{23} = -1, a_{24} = -1$. The dna_dsb input vector is held at a constant value 1, meaning that the system is constantly under DNA damage stress. We assume in this experiment negative regulation biases, $b_i = -1/2$, for $i = 1, 2, 3, 4$. The transition noise parameter $p$ is selected randomly in the interval $[0.01, 0.1]$. The microarray data model has parameters $\mu_i \equiv \mu = 30$, $\delta_i \equiv \delta = 20$, $\sigma_i^2 \equiv \sigma^2 = 49$, for $i = 1, \ldots, 4$. Average edge-calling error rates obtained by PALLAS over 20 repetitions of the experiment are displayed as a function of time series length in Figure 4. One can see that, as the time series length increases, precision, sensitivity and specificity all increase. It can be seen also that performance improves quickly initially, but after the time series length exceeds 20 there is little additional improvement.

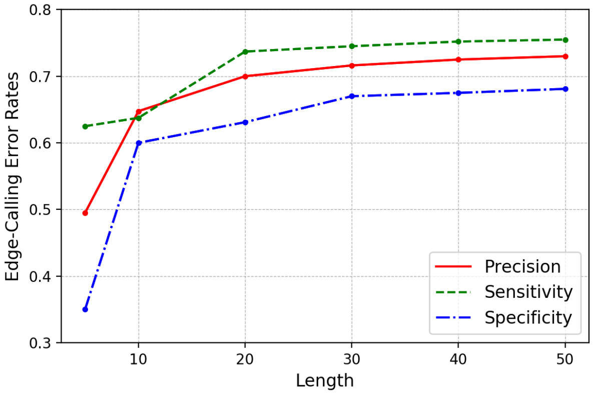


Figure 4: p53-MDM2 experiment edge-calling error rate results as a function of time series.

### 4 E. Coli SOS DNA Repair System

Figure 5 displays the full network obtained by PALLAS as a consensus of the three top networks according to penalized likelihood score.

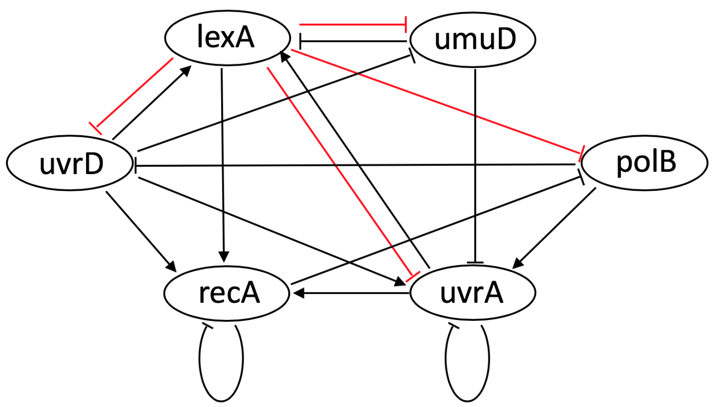


Figure 5: The SOS DNA repair system network inferred by PALLAS

### 5 E. Coli Biofilm Architecture

Figure 6 displays the full network obtained by PALLAS as a consensus of the three top networks according to penalized likelihood score.

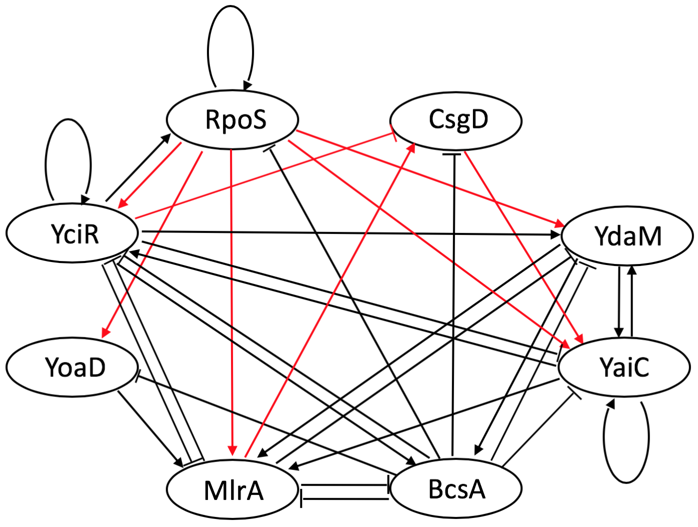


Figure 6: The biofilm system network inferred by PALLAS
